# Periodic Non-Sinusoidal Activity Can Produce Cross-Frequency Coupling in Cortical Signals in the Absence of Functional Interaction Between Neural Sources

**DOI:** 10.1101/062190

**Authors:** Edden M. Gerber, Boaz Sadeh, Andrew Ward, Robert T. Knight, Leon Y. Deouell

## Abstract

The analysis of cross-frequency coupling (CFC) has become popular in studies involving intracranial and scalp EEG recordings in humans. It has been argued that some cases where CFC is mathematically present may not reflect an interaction of two distinct yet functionally coupled neural sources with different frequencies. Here we provide two empirical examples from intracranial recordings where CFC can be shown to be driven by the shape of a periodic waveform rather than by a functional interaction between distinct sources. Using simulations, we also present a generalized and realistic scenario where such coupling may arise. This scenario, which we term waveform-dependent CFC, arises when sharp waveforms (e.g., cortical potentials) occur in a periodic manner throughout parts of the data. Since the waveforms are repeated periodically, they constitute a slow wave that is inherently phase-aligned with the high-frequency component carried by the same waveforms. We submit that such behavior of the data, which seems to be present in various cortical signals, cannot be interpreted as reflecting functional modulation between distinct neural sources without additional evidence. In addition, we show that even low amplitude periodic potentials that cannot be readily observed or controlled for, are sufficient for significant CFC to occur.

## Introduction

Phase-amplitude cross-frequency coupling (CFC) refers to a dependence between the phase of a slow frequency ("frequency-for-phase") and the power of higher-frequency activity ("frequency-for-amplitude") recorded from the brain (1–3). Numerous studies have provided evidence that it may play an important role in behavior and cognition (4–9; for review, see 2,10,11). CFC is usually interpreted in the context of two distinct processes that are coupled such that the slow frequency component of one process drives, or modulates, the high frequency component of the other. A mathematical dependence between two frequency bands, on the other hand, is not by itself a sufficient indication of such an interaction (1,12). A recent theoretical account by Aru et al. posits that any non-stationary process in the signal can affect both the phase of a low-frequency component and the amplitude of a high-frequency component, thus generating spectral dependencies which would be interpreted as CFC, even though they are driven by a single source (12). Using simulated data, Kramer and colleagues argued that under certain circumstances sharp edges in the data may result in coupling in a wide range of frequencies-for-amplitude (13). However, this type of coupling has not yet been demonstrated in real scalp-or intracranial-EEG data, and it remains unclear under what circumstances, if at all, such a situation may arise. In this report we present real-world examples of intracranial EEG recordings where CFC is likely to be caused by the temporal characteristics of a single process rather than by an interaction between two processes, and demonstrate how such a scenario can realistically be manifested in any EEG signal.

Specifically, if a sequence of sharp periodic waveforms (even if jittered or intermittent) exists in the data (e.g. due to neuronal potentials, electrical interference, electrocardiographic potentials, etc.), it may manifest as strong CFC. This happens because the periodicity of non-zero-mean sharp deflections constitutes a low frequency component, which is inherently coupled with the amplitude peaks of the high frequencies contained in the sharp deflections themselves, in the absence of any other causal mechanism driving this dependence. When the data are filtered to measure coupling, the phase at the frequency of occurrence of the potentials will align with their peaks, introducing robust CFC (waveform-dependent CFC). Additionally, if the occurrence of the periodic potentials persists sufficiently in time, even very low amplitude waveforms may suffice to introduce significant CFC. We demonstrate the above claims using a series of simulations, and show evidence from electrocorticography (ECoG) data that such a scenario occurs in certain commonly observed oscillatory signals, such as the mu rhythm (8-10Hz) and the beta rhythm (13-20Hz) in sensorimotor areas, which often assume the shape of a sequence of sharp deflections rather than smooth oscillations (Fig. 4).

## Materials and Methods

*Simulation of waveform-dependent CFC*. We first demonstrate how a semi-periodic occurrence of sharp waveforms can drive waveform-dependent CFC in EEG data, by simulating trains of Gaussian-shaped spikes of parametrically varied heights and widths and peak-to-peak intervals (frequency of occurrence) and introducing them into otherwise non-coupled signal traces. The height of the Gaussian spikes is given in units of standard deviations (STDs) of the background signal, and set to either 1.5 or 3 STDs. The width of the Gaussians is given as the full-width-at-half-maximum, and set to 10 or 20 milliseconds (sampling rate: 1000Hz). The spikes occurred at one of two jittered intervals: either at a mean interval of 100±20ms peak-to-peak (uniformly distributed), corresponding to a 10Hz spectral peak with a bandwidth of about 3Hz, or at a mean interval of 167±33ms corresponding to a 6Hz spectral peak with a bandwidth of about 2.5Hz. These peak-to-peak time interval values were selected to mimic coupling with alpha and theta as the frequencies-for-phase, respectively. The fairly large variability around this interval, of about 20% of the mean interval, were used in order to give the simulation a physiologically realistic nature and to make it similar to the empirical cases shown later.

Each spike train was superimposed on to a 60-second trace of background activity, consisting of either a 1/f pink-noise simulated signal, or real EEG channel data. A 60-second trace of pink noise with a sampling rate of 1000Hz was generated using the method outlined by Kasdin (1995) (14); briefly, we generated random white noise and then applied a series of filters in a pseudo-continuous manner to produce a 1/f distribution at our specified sampling rate. The scalp EEG data used as background activity consisted of 60 seconds of continuous scalp EEG, recorded from a healthy subject performing a visual perception paradigm designed for other experimental purposes. The occipital electrode Oz was chosen from a subject who showed no CFC. Data were recorded with a 64 channel BioSemi EEG system at 1024Hz sampling rate. After recording data were resampled to 1000Hz, re-referenced to the common average, notch filtered at 60Hz, and high-pass filtered at 0.5Hz using the two-way, zero phase-lag, FIR filter implemented in the *eegfilt* function of the EEGLAB toolbox for Matlab (15). This EEG dataset features the common ~1/f property of its spectral content, but also has a local power peak around the alpha band, as it was taken from an occipital electrode. By using real data as one of the two base signals for our simulations we wanted to assure the ecological validity of our tests, and to examine whether the artificially added periodic activity at 10Hz generates CFC in data with a preexisting prominent physiological alpha oscillation. The total number of simulated traces was therefore 16 (2 Gaussian heights × 2 Gaussian widths × 2 inter-peak intervals × 2 background signals). One of the aforementioned simulated traces (amplitude: 3 STDs; width: 10 ms; mean interval: 100 ms, superimposed on the scalp EEG signal) was further used for comparison with the empirical data cases (see Figure 4C).

*Simulation of coupled-sources CFC*. To simulate an alternative case of physiologically realistic coupling, reflecting modulation of gamma power by the phase of an oscillatory alpha rhythm rather than a periodic series of sharp waveforms, we extracted the 30-100Hz band from the EEG signal, and multiplied it point-by-point by a phase index factor. This factor is simply a vector of values ranging continuously between 0.75 at the troughs of the 10Hz cycle of the signal and 1.25 at the peaks of the signal’s 10Hz cycle. When the extracted gamma was multiplied by this factor, the gamma power was forced into a correlation with the alpha phase. We then subtracted the original 30-100Hz band and inserted the modulated one in its place. This manipulation effectively produced robust coupling between the amplitude of the broadband gamma power and the phase of the alpha band, which cannot be attributed to a consistent waveform shape.

*Real data cases - electrocorticography (ECoG)*. We report two cases of waveform-dependent CFC in intracranial electroencephalographic recordings. ECoG recordings were made in patients implanted with subdural electrodes for the treatment of intractable epilepsy at the Stanford School of Medicine, as part of unrelated ongoing studies. The electrocorticographic recording exhibiting mu-oscillations were obtained from a patient performing a task which involved viewing images from several categories and responding by a button press to a target category. The electrode included in this study was located on the right primary somatosensory cortex of the patient. The recording exhibiting beta oscillations was obtained from a patient performing a picture naming task in which she was required to name line drawings of various objects appearing on the screen. The electrode included in this study was located on the right primary motor cortex. Subjects gave informed consent approved by the UC Berkeley Committee on Human Research and corresponding IRBs at the clinical recording site. Both datasets were recorded with a Tucker-Davis Technologies (TDT) recording system at a sampling rate of 3051.76Hz. After acquisition, data were re-referenced to the common average, resampled to 1000Hz, and high-pass filtered at 0.5Hz. The electrodes selected for this study did not show epileptic activity.

*Simulation of low-amplitude waveforms in EEG data*. In order to examine whether sharp periodic waveforms can drive CFC in realistic EEG settings where they may be of considerably low amplitude (relative to background activity), we extended our previous simulations of periodic activity. Using the scalp EEG recording as model of background EEG activity and a 100±20 ms spike interval, we examined the magnitude of the resulting CFC for a range of smaller spike amplitudes as well as several widths. Amplitudes ranged from 0.4 to 1.8 standard deviations of the background signal in 0.1 steps, while full-width at half-maximum values ranged from 1 to 25 ms in single-millisecond steps. CFC indices and statistical significance were subsequently computed for each width/amplitude combination using 10Hz as frequency-for phase and the optimal frequency-for-amplitude for the relevant spike width (see statistical analysis below).

*Phase-amplitude cross-frequency coupling analysis*. We assessed the magnitude of coupling between high frequency power and low frequency phase using the phase-locking value method (PLV) (11,16–19). In short, for each combination of frequencies, two new time series were created from the raw signal by filtering it in the frequency for phase (f-p) and the frequency for amplitude (f-a) bands. The Hilbert transform was then applied to each one, from which we extract the instantaneous phase of f-p as well as the amplitude envelope of f-a. In order to evaluate the PLV between the two, both should be expressed as instantaneous phases. To that end the amplitude envelope signal is filtered at the frequency values used to create f-p, and the phase information is extracted from the filtered signal via a second Hilbert transform. The PLV is then computed between the two phase series, and ranges between 0 (no phase-to-amplitude relation) to 1 (perfect phase-to-amplitude relation).

For the frequency-for-phase component, signals were band-pass filtered using a 2Hz band around the center frequency. The band-pass filter for the frequency-for-amplitude was adaptively determined based on the frequency-for-phase, since the amplitude modulation of a signal produces spectral side-bands equal in bandwidth to the modulating signal (12). For instance, phase-amplitude coupling between a 10Hz and a 60Hz tone produces sidebands at 50 and 70Hz. Neglecting to include this entire bandwidth when filtering the frequency-for-amplitude signal prevents the full extent of CFC to be measured, leading to false-negative results. We therefore set the bandwidth of each filtered frequency-for-amplitude component based on the corresponding frequency-for-phase (e.g., for a 20Hz frequency-for-phase, we applied a passband of 40Hz around the frequency-for-amplitude).

*Statistical analysis*. As a preliminary step, we identified the preferred frequency-for-amplitude for which CFC occurs for each Gaussian width value. This was done so that a single optimal CFC measure can be produced for a given signal. For this purpose, we injected a 10Hz spike train with a fixed amplitude of 5 standard deviations to a background signal to produce robust CFC, and the magnitude of CFC was determined for a 1-200Hz range of frequencies-for-amplitude, with 10Hz being fixed as the frequency-for-phase (the periodic potentials produced a clear spectral peak at this frequency). The preferred frequency-for-amplitude was selected as the one producing the strongest CFC. The results were averaged over 100 repetitions of this procedure to increase their accuracy.

Statistical significance was assessed using a bootstrap procedure. For each frequency pair, one of the two filtered time series (frequency-for-amplitude) was circularly shifted in time by a random offset, and the CFC index was calculated for the resulting misaligned traces (12,20). This operation was repeated 2000 times to generate a distribution of surrogate modulation indices for each frequency combination. Statistical significance was calculated as the proportion of surrogate values smaller than the original CFC index, using the equation p = (r+1)/(n+1), where r is the number of surrogates higher than the observed statistic, and n is the total number of surrogates (21).

*Phase-phase coupling analysis*. As we discuss below, the periodic repetition of sharp potentials may also lead to a phase-phase correlation between the low and high frequency components of the resulting CFC. This is because the train of potentials drives the phase of both the slow and high frequency components of the CFC. We therefore wanted to test whether phase-phase relations could constitute a means of detecting the presence of periodic sharp potentials driving a CFC effect. To observe phase-phase dependencies, the phase data of both the low-frequency and high-frequency components were extracted using the Hilbert transform after band-pass filtering the signal with a bandwidth of 2Hz. The instantaneous phases of each cycle were then binned into 20 bins and a histogram was produced for the phase bins of the high-frequency signal co-occurring with the preferred low-frequency phase of the CFC (i.e. the low-frequency phase at which high-frequency amplitude is maximal). A non-phase locked interaction would entail a random, uniform distribution of high-frequency phases, while any other distribution would indicate a dependency between the phase signals. The coupling was assessed quantitatively using the Rayleigh test, which provides a significance level for the length of the vector produced by the complex mean of the phases on the unit circle. A significant result reflects a nonrandom phase concentration around a specific value. The Rayleigh test was implemented using the Circular Statistics Toolbox (22). All processing was achieved using custom routines in MATLAB (MathWorks Inc.).

## Results

*Simulating waveform-dependent CFC in realistic EEG models*. To examine how phase-amplitude coupling is generated in signals merely through the introduction of semi-periodic potentials, we generated 8 Gaussian spike trains corresponding to combinations of waveform amplitude (1.5 vs. 3 STDs), width (10 vs. 20 ms), and mean inter-spike interval (100 vs. 166 ms, corresponding to a 10 vs. 6Hz periodicity), and superimposed each on both a scalp EEG and a simulated pink-noise background signal, totaling in 16 traces (see Methods). The background time series alone did not result in significant clusters of coupling at either the alpha or theta simulated frequencies-for-phase (nor in other frequencies, Figure 1A,E). In contrast, as expected, all compound signals did feature strong CFC, as evidenced by the robust clusters of statistically significant coupling in the comodulograms in Figure 1B-D and F-H (all p-values < 0.005 at the preferred frequency-pairs). Note that for the sake of space Figure 1 includes visualization of 6 out of the total 16 simulated cases, however all 16 compound traces, with either EEG or pink noise as background signals, resulted in significant clusters of coupled frequency pairs.

**Figure 1.**
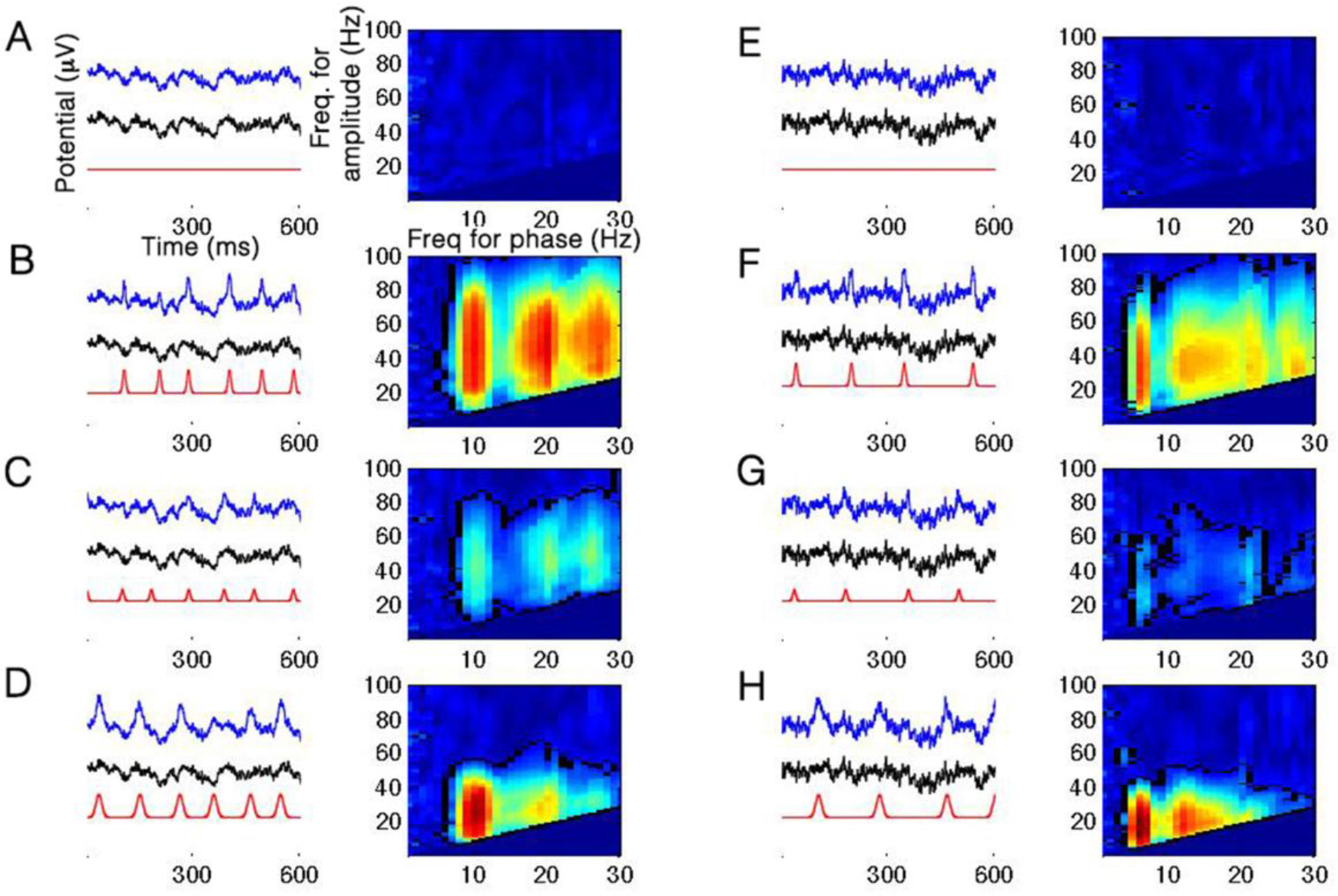
Examples of CFC in six simulated signals. The plots on the left of each column depict a trace from the original (background) signal used for the simulation (black), the added Gaussian train (red), and the compound signal after adding the potentials (blue). The blue trace is therefore the sum of the black and the red ones. Note the temporal jitter in the Gaussian periodicity (i.e. variability of the inter-peak interval). The image on the right in each column shows the comodulogram for the compound signal (blue to red color scale corresponds to 0 to 1 in all panels). Clusters of significant CFC in the comodulograms are marked with a black outline (p<0.01). Note that values in the bottom triangle of each plot are not shown as CFC is invalid for frequencies-for-amplitude lower than the frequency-for-phase. The left column shows results for EEG-based simulations with a 100-ms mean inter-peak interval; the right column shows results for pink-noise-based simulations with a 166-ms mean inter-peak interval. It can be easily seen that the periodicity determines the frequency-for-phase of the CFC (10Hz and 6Hz, respectively), as well as its harmonics. A, D, the raw background traces with no added spikes. B, F, added spikes with an amplitude of 3 STDs and a width of 10 ms (full-width at half-maximum). C, G, Same as B,F but with an amplitude of 1.5 STDs. Note the diminished magnitude of the effect. D, H, Same as B,F but with a width of 20 ms. Note the lower frequency-for-amplitude range for which CFC occurs.

Inspection of the right vs. left column of Figure 1 clearly indicates that the CFC effect is not contingent on a specific frequency band. The frequency-for-phase reflects the mean peak-to-peak interval of the Gaussian spike trains such that the 100±20ms trains predictably produced significant CFC around frequency-for-phase of 10Hz, and the 167±33ms trains similarly produced significant CFC around frequency-for-phase of 6Hz. The comodulograms also feature gradually decreasing CFC at subsequent harmonics of this frequency. Secondly, the amplitude of the simulated Gaussians influences the coupling such that larger potentials lead to stronger PLV values (Figure 1B,F vs. C,G). Lastly, comparing Figure 1B,F with D,H shows how the width of the waveform determines the range of frequencies-for-amplitude, such that narrower, or more “spiky”, activity produces CFC in a higher range of frequencies-for-amplitude, as narrower waveforms correspond to higher frequency content. This relation is shown in detail in Figure 2, which plots the frequency-for-amplitude corresponding to the strongest CFC as a function of spike width (see Methods).

**Figure 2.**
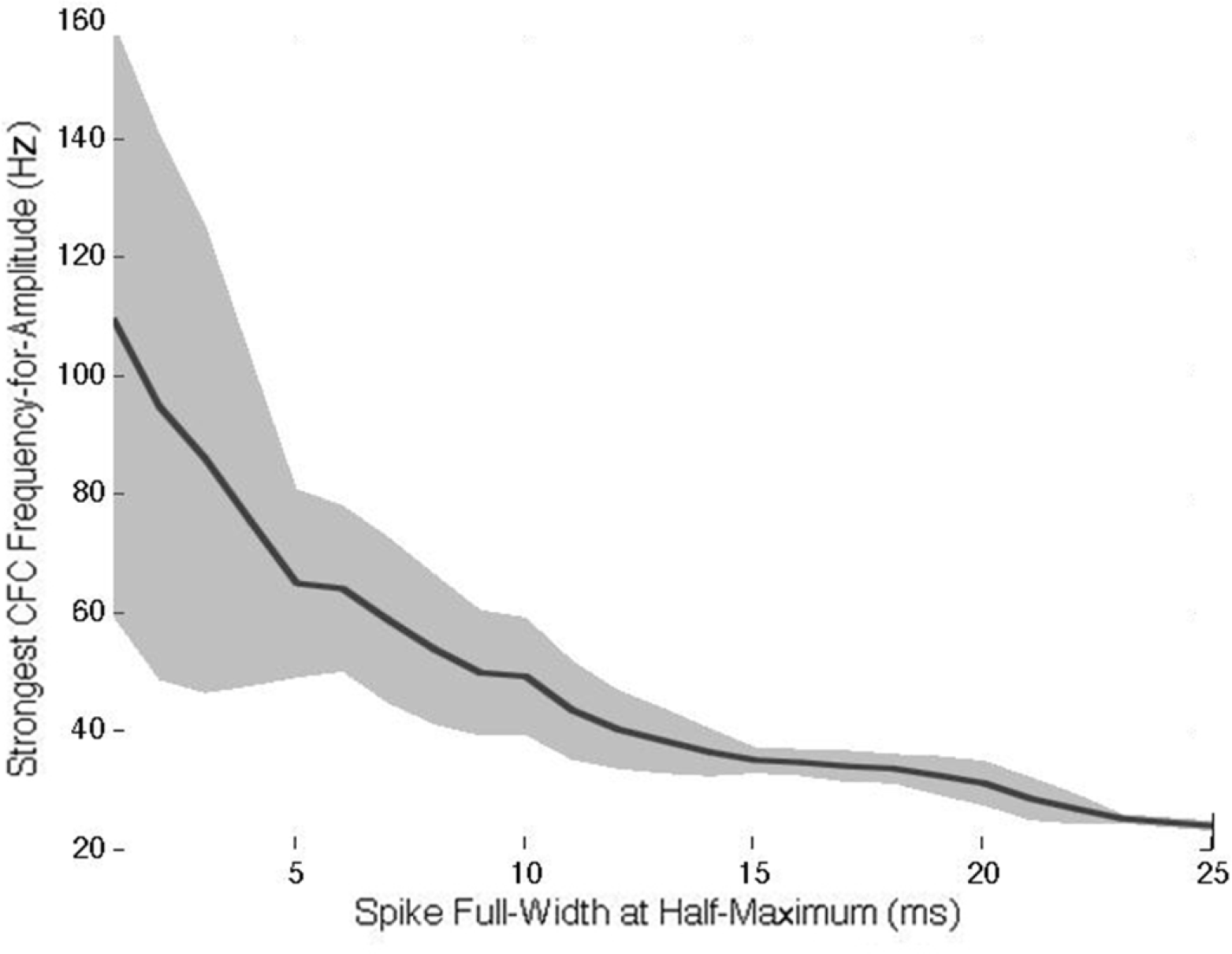
Preferred frequency-for-amplitude as a function of Gaussian spike width. Shaded area reflects one standard deviation of the preferred frequency across 100 iterations (see Materials and Methods).

Phase-amplitude coupling in these cases occurs since the same recurring waveforms drive both the phase of the low-frequency component (i.e., narrowly filtering the signal around 10Hz generates an oscillation whose peaks correspond to the location of the spikes) – as well as the amplitude envelope of the high-frequency component (since each sharp peak corresponds to a local increase in high-frequency power). Unlike in the specific case described by Kramer (2008) (13), this effect does not depend on the high-frequency potentials occurring systematically on specific phases of an existing low-frequency oscillation, since here the recurring waveforms themselves define the low-frequency oscillation. To clarify this, we examine what happens to the phase of the slow oscillation (the frequency-for-phase of the CFC) contained in the background signal following the addition of the spike trains: Figure 3 shows what happens when a series of 10Hz periodic potentials with a magnitude of 1.5STDs of the background scalp EEG signal is injected into the background signal which already features a prominent 10Hz oscillation due to physiological alpha rhythms recorded over the occipital scalp. The peaks of the potentials were inserted at random phases of the 10Hz oscillation of the original signal (blue bars). However, after their addition to the signal, the 10Hz 0-phase (peak) values of the resulting compound signal tended to concentrate at the locations where the spikes had been inserted (Figure 3A). That is, the added spikes caused a shift in the phase of the 10Hz frequency component such that the peak phase tends to align with the peak of the potentials (Figure 3B), despite not significantly affecting the spectral magnitude of this component (Figure 3C). Note that the spikes were mathematically added to the signal after it was recorded, and thus could not actually affect the physiological processes eliciting the alpha rhythm (i.e., no true entrainment is involved)

**Figure 3.**
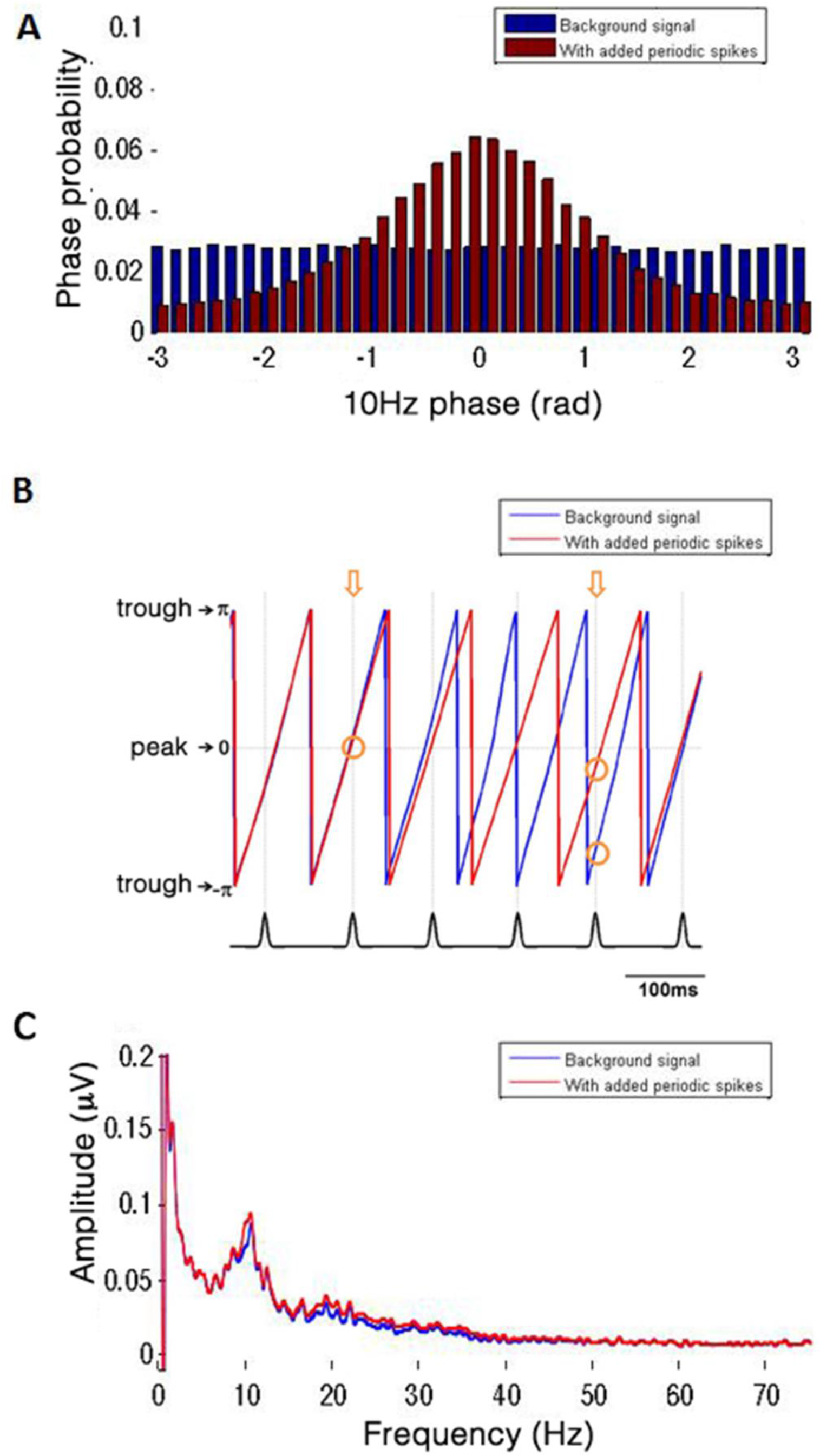
Phase-shift of the background EEG signal after the addition of periodic potentials. The signal was generated by adding a Gaussian spike train with 100-ms mean inter-peak interval, 15-ms width and amplitude of 1.5 STDs to the background EEG trace. A, Distribution of 10Hz phase values at the time points where Gaussian peaks occurred. Blue: 10Hz phase extracted from the background EEG signal before adding the Gaussians train; red: phase extracted from the compound signal. It can be seen that while the spikes were inserted at random points of the 10Hz cycle of the background signal, the same time points show a prominent phase-concentration around the cycle peak (phase 0) once the Gaussians were integrated with the signal. That is, the 10Hz cycle tends to align with the transient periodicity such that a phase-amplitude relation emerges. B, Example of alpha phase change of the EEG background signal in a few consecutive cycles, before and after the introduction of the spike train. Blue trace represents the original signal, red trace represents the compound signal. Note that as the Gaussian peak falls away from the original alpha peak, the new alpha cycle will shift farther in phase (compare the two arrows). C, This rearrangement of the alpha phase can occur without any significant change to the power spectrum.

*Waveform-dependent CFC in intracranial mu and beta signals*. We report two cases of suggested waveform-dependent CFC in intracranial signals, in the mu and beta rhythms. Mu oscillations, which are typical in sensory motor areas, are characterized by periodical sharp deflections rather than smooth sinusoidal oscillations (anecdotally, the name “mu” was attributed to this activity because of the similarity between the shape of the oscillatory cycle and the Greek letter μ (23,24)). Similarly, previous reports of beta oscillations in the motor cortex (also called Rolandic beta) show a similar spiky appearance of the oscillation (25–27). Considering the above simulations, this characteristic of sharp periodic waveforms might produce significant phase-amplitude coupling, in the absence of an interaction between two distinct neural processes operating at different frequencies. Under this scenario, the periodic repetition of high-amplitude sharp potentials would drive both the phase of the low-frequency and the periodic increase in high-frequency power, which would therefore be intrinsically coupled.

The recorded Mu and Beta rhythms (Figure 4A and 4B, respectively) are characterized by a sharp, high-frequency waveform occurring periodically, resulting in a strong, statistically-significant CFC between the corresponding frequency phases (at ~10 and ~17 Hz respectively) and a broad range of gamma frequencies. Note that similarly to our simulations of waveform-dependent CFC, both the Mu and the Beta signals feature CFC at the first subsequent harmonic of the frequency-for-phase, corresponding to the harmonics generated by a non-sinusoidal sharp periodic waveform (For the mu signal, phase-amplitude coupling is significant at p<0.01 between the first harmonic of 20Hz as frequency-for-phase and the 22-78Hz band of frequencies-for-amplitude; likewise for the beta signal, coupling is significant at p<0.01 between 34Hz and 36-60Hz). For comparison purposes we present the ECOG data along with two cases of simulated signals from our simulations above – a train of sharp periodic potentials (3STDs height, 10ms width, 100ms (±20ms) inter-peak interval) superimposed on the background EEG signal (Figure 4C), and a simulated coupled-sources CFC produced by explicitly modulating the high-frequency (30-100Hz) component of the raw EEG signal by the phase of its 9-11 Hz oscillation (see Methods; Figure 4D). The introduction of the train of sharp potentials elicits a CFC a the fundamental frequencies and its harmonics. The simulated coupled-sources CFC (Figure 4D), on the other hand, represents a strong coupling without harmonics owing to the smooth nature of the low-frequency periodicity.

**Figure 4.**
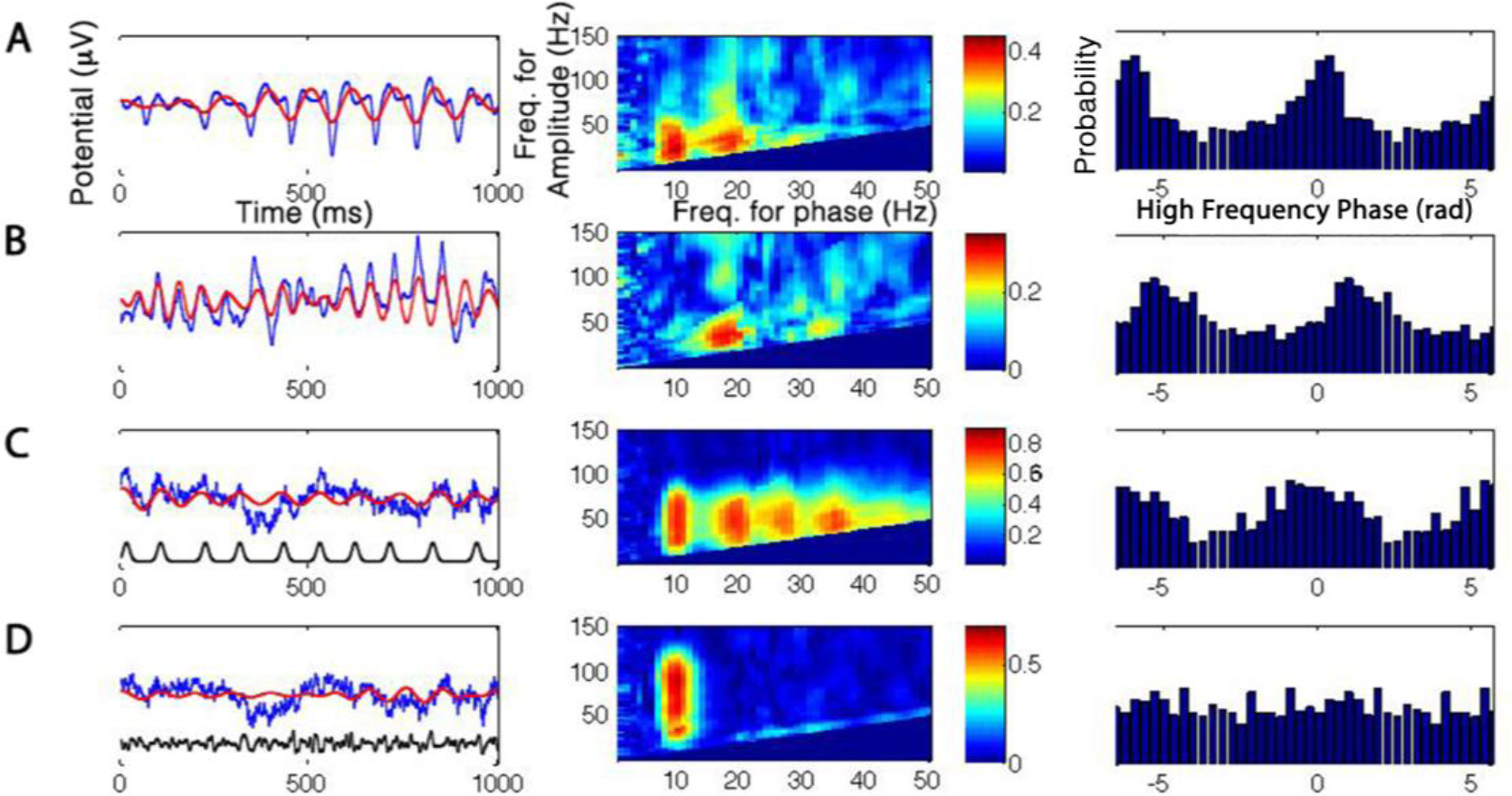
Waveform-dependent CFC in empiric signals. Left column: example traces of each analyzed signal. The blue trace is the raw trace, and the red is the low-frequency band-pass filtered component. Black trace in the bottom two rows shows the artificial data added to induce CFC. Middle column: comodulogram showing the PLV value for each frequency-pair. Right column: Phase-phase dependency: histogram of high-frequency phase values (in radians) occurring at the preferred phase of the low-frequency cycle (i.e. the phase for which high-frequency amplitude is maximal) for the basic rhythm frequency (10Hz for A,C,D, 17Hz for B) with its first harmonic (20Hz for A,C,D, 34Hz for B). The histogram is repeated twice along its horizontal axis to improve readability. A, Intracranial recording of mu oscillations (~10Hz). B, Intracranial recording of beta oscillations (~17Hz). Note the non-sinusoidal single sharp peaks driving the oscillatory component in A and B, as well as the harmonics in the comodulogram and the non-uniform phase-phase relation between the low-and high-frequency components. C, CFC induced by superimposing a periodic Gaussian train on a background EEG trace, featuring no CFC initially. D, "Coupled-sources" CFC induced by modulating the amplitude of the 30-100Hz band by the instantaneous 10Hz phase. Note the similar behavior of A, B compared to C and not to D.

To further test our claim, we proceeded to examine the phase-phase relations between the coupled frequencies in each case. Waveform-dependent CFC entails coupling between the phases of the two frequencies, as the peaks of the periodic waveforms drive the phases of both frequency components. In contrast, the common interpretation of phase-amplitude coupling, i.e. the mechanistic interaction of two sources of neural activity whereby the phase of one modulates the magnitude of the other, does not entail any statistical dependence between the instantaneous phases of the paired frequencies. While phase-phase relations may in principle occur in cases of such an interaction between sources, it is a very specific case and calls for care in ruling out simpler alternative explanations. The right column of Figure 4 demonstrates this distinction. Phase-phase coupling was assessed in the simulated and empirical cases by plotting a histogram of the high-frequency phases cooccurring with the preferred low-frequency phase of the CFC interaction. When no phase-phase dependence is expected, as in the second simulated case (Fig 4D), the probability of any phase-pair to occur is equal, leading to a uniform distribution. However in the case of CFC driven by periodic spikes, the phase-resetting of multiple frequencies around the peaks leads to a non-uniform phase distribution (Fig. 4C). The empirical cases (4A and 4B) are similar to the spike-train scenario in showing a non-uniform distribution. This lends further evidence to the claim that CFC in these cases can be parsimoniously explained as resulting from a series of sharp potentials rather than from the modulation of a high-frequency source by the phase of a low-frequency source.

*Simulating low amplitude spike trains*. The existence of waveform-dependent CFC in ECoG recordings serves as a clear indication that CFC may reflect the temporal characteristic of non-sinusoidal periodic waveforms. To test whether the presence of CFC in the absence of non-sinusoidal periodic waveforms in the data could be used as an argument for functional coupling between two sources, we extended our simulations to test the effect of low-SNR periodic potentials in a realistic EEG signal. There are various sources that could potentially contribute such low SNR activity to a recorded signal, whether external or biological (e.g. heartbeat, muscle activity, induced rhythmic potentials, event-related or other cortical potentials, etc.).

We generated a semi-periodic spike train of 100±20ms intervals for each width and amplitude parameter combination detailed above, injected it onto the background signal, and statistically tested for phase-amplitude coupling between the fixed frequency-for-phase (10Hz), and the preferred frequency-for-amplitude previously determined for the current spike width (see Methods). This was repeated 100 times to increase the accuracy of the results, presented in Figure 5A. Instead of presenting the raw PLV, we present the degree to which the results were statistically significant. Our reasoning is that in a common scenario where a signal is analyzed for CFC, typically several frequency-pairs will be included in the analysis and the results appropriately corrected for multiple comparisons. Assuming a basic type I error probability rate of 0.05, Figure 5A plots the number of multiple comparisons that the test would pass according to the mean p-value for each parameter combination (width and amplitude) at the preferred frequency pair (computed as *n* = 0.05/*p_value*), resulting in a simple linear scale where stronger CFC corresponds to a higher significance value (i.e. number of comparisons). The results show that CFC may be generated even when the amplitude of the periodic potentials is no more than one standard deviation of the background signal. In this kind of scenario there would be no clear indication, especially by visual examination of the data, that the CFC effect is driven by periodic spikes (see raw data trace in figure 4C, where spike amplitude is set at 1.5 STD) unless this hypothesis is explicitly investigated. Furthermore, harmonics in the comodulogram are of much lower intensity for low-amplitude spikes (compare, for example, the results for 1.5STD spikes and 3STD spikes in figure 1), making this indication for non-sinusoidal periodicity unreliable as well. Such a scenario may be realized, for example, in scalp EEG recordings or in ECoG electrodes remote from the source, where the spiky nature of mu, beta, or any other oscillatory activity may be less distinctively discernable due to lower SNR and a greater spatial summation of sources.

**Figure 5.**
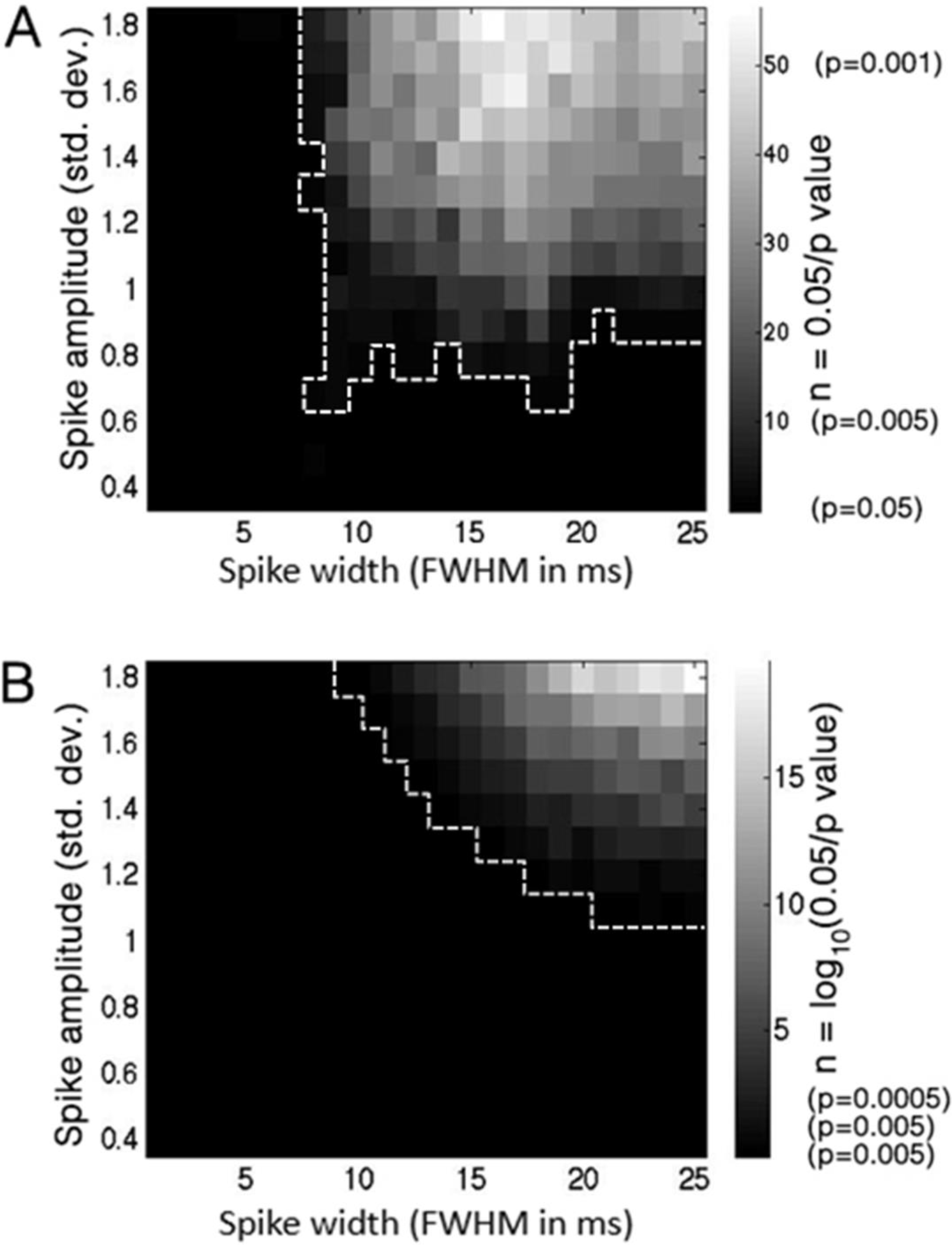
Statistical significance of phase-amplitude and phase-phase coupling for low-amplitude transients. A, Each pixel in the image represents the average p-value, across 100 repetitions, for a phase-amplitude CFC effect driven by adding a Gaussian spike train (inter-spike interval: 100 ms) of the given amplitude and width, to a background EEG signal featuring no initial CFC. The colorscale corresponds not to the actual p-value, but to n=0.05/p, i.e. the number of Bonferroni multiple-comparisons the effect “survives”, resulting in a simple linear scale of significance. The white outline indicates the cluster of significant pixels (n>=1). Importantly, note that significant CFC occurs even for very low-amplitude transients (under 1 standard deviation). It can also be seen that the magnitude of CFC depends on the width of the transient waveform: very narrow transients do not contain enough energy to drive the phase of the slow-frequency component, while wide transients contain a weaker high-frequency component. B, Significance of phase-phase coupling between 10Hz and 20Hz is shown for the same range of transient amplitudes and widths. Since a parametric significance test was used, p-value tended to be very small and so the logarithmic scale n=log10(0.05/p) was used (such that n=0 corresponds to p=0.05, n=1 to p=0.005, etc.). White border indicates area where p<0.05. Note that the area indicated as significant in this analysis does not fully overlap that of the phase-amplitude coupling analysis (Figure 5A).

We showed above that spectral coupling driven by sharp semi-periodic activity can give rise to strong phase-phase relations between the frequency-for-phase and the frequency-for-amplitude (Figure 4, right column). To test the possibility that phase-phase correlation could be a sensitive test for the waveform-driven nature of a CFC effect generated by small amplitude sharp potentials, we ran a phase-phase coupling analysis on the same array of simulated signals as for the phase-amplitude analysis above, producing a comparable matrix of phase-phase-coupling significance values (figure 5B). Phase-phase coupling was assessed in each case between the 10 and 20Hz components, as the first harmonic frequency was consistently found to produce the most robust phase-phase coupling for any given lower frequency-for-phase in the case of periodic potentials (note that this is in contrast to the case of phase-amplitude coupling, where the optimal upper-frequency is determined by the waveform characteristics as described earlier). The results show that while highly-significant phase-phase coupling is produced for high-power spike trains (amplitude × width), there is a considerable range of cases where significant phase-amplitude coupling is produced by low-amplitude potentials while no significant phase-phase coupling can be detected (compare Figure 5A and 5B). We conclude that the absence of phase-phase coupling is not always a reliable indicator for the absence of low-amplitude waveform-dependent phase-amplitude coupling.

## Discussion

Phase-amplitude cross frequency coupling is commonly taken as a signature of functional coupling between low and high frequency signal sources. However, a periodic repetition of non-sinusoidal waveforms, such as cortical potentials or external artifacts, also leads to the same spectral signature due to the alignment of the phase of the slow periodicity with the high-frequency power fluctuations driven by the same waveforms. In this case, the CFC merely reflects the shape of the waveforms and the fact they occur with regular intervals. In this report we show examples from ECoG data in which observed CFC may result from sharp periodic potentials, driving inherently-coupled high frequency power and low frequency phase in the absence of any functional modulation. The simulations used to demonstrate this principle also show that as can be expected, the shape of the waveforms determines the range of frequencies-for-amplitude, their amplitude determines the strength of the resulting coupling, and their rate of occurrence determines the frequency-for-phase.

Artifacts resulting in spurious coupling were previously studied (12,13). In their report, Kramer and colleagues convincingly showed how ripples created around edges and voltage steps that exist in the signal can generate coupling in a wide range of frequencies-for-amplitude. However, in these simulations the voltage steps were inserted around the same phase across cycles of the slow oscillation, and the demonstration was centered on the high frequency properties of the signal. Although of high theoretical importance, the extent and the context in which such confounding factors appear in real data are unclear. The cases presented in the present report are realistic in the sense that any activity, physiological or not, characterized by sharp potentials appearing in a roughly periodic manner (even if fairly jittered), is enough to produce CFC. The demonstration of waveform-dependent CFC resulting from common electrophysiological activity like mu or beta emphasizes the challenge in interpreting CFC results in EEG and ECOG data in a physiologically meaningful way.

Periodic sharp waves are reflected in CFC analysis as bursts of high frequency power occurring at the peaks of a slow oscillation. While this can also be a signature of a true dual process CFC, whereby a source of sharp potential is coupled to a slow oscillation at the peak of its cycle, such interpretation is less parsimonious than waveform-dependent CFC (implying a single process) and requires additional evidence. One can also argue that the existence of periodic sharp waves *per se* implies, by necessity, a low-frequency process (pacemaker) which triggers the individual spikes. Similarly, any oscillation which is not strictly sinusoidal will be reflected spectrally as periodic epochs of higher frequency activity. However, for CFC analysis to be applied meaningfully, it should attempt to differentiate cases where the data supports functionally-coupled sources from cases of simple non-sinusoidal periodicity. An example for the former case could be coupling between a theta-band oscillatory LFP signal and a higher-frequency, zero-mean activity generated by spiking. In this case, the spiking activity could hypothetically occur on any part of the theta oscillation cycle without altering the latter's phase, and CFC is assumed since the spikes nevertheless consistently occur on a specific phase; in contrast, in the mu/beta signals above the occurrence of the high-frequency spikes could not be shifted in time without also shifting the low-frequency phase component, thus maintaining the waveform-dependent phase-amplitude coupling.

The above arguments refer to the interpretation of the low frequency component (phase component) of the CFC. Similar attention is due to the high frequency component. While the basic definition of phase-amplitude-coupling makes no assumptions as to the signal characteristics driving the high-frequency amplitude, meaningful interpretation requires specification of the type of waveforms responsible for this component. The amplitude of a given frequency band could be driven by a band-limited oscillation, but also by wideband noise or by individual sharp potentials (as is the case shown here). While we do not argue that the concept of CFC should only be reserved for a specific case, it is vital for the understanding of neural phenomena that we distinguish between such different manifestations of spectral modulations. Particularly, care should be taken not to imply an interaction between slow-frequency and high-frequency oscillatory processes where the coupling is in fact driven by wideband phenomena.

Our simulations suggest that there is no single criterion that can distinguish waveform-dependent CFC from other mechanisms. However, several characteristics of the signal are suggestive of waveform-dependent CFC:

1. Phase-phase coupling was a characteristic for the mu and beta cases, as it was for simulated cases where the magnitude of the spike trains was large enough to induce precise phase-locking of the frequency components. However, we also show that this test may not be sensitive enough when the potentials are of low amplitude, whereas they may still be large enough to result in significant phase-amplitude cross-frequency-coupling
2. Harmonics in the frequency spectrum and in the comodulogram may be an indication that the raw signal contains non-sinusoidal periodic waveforms that may contain higher-frequency components, as evident in Figures 1 and 4.
3. The raw data should be visually inspected in an effort to identify the source of the coupling. Overlaying the band-pass filtered slow-wave component over the raw data trace (e.g. as in Figure 4, left column) can provide some indication of when the rhythmicity of the signal consistently deviates from a smooth oscillation, raising the possibility of a periodic waveform with a high frequency component.
4. A strong argument in favor of a two-process interaction interpretation of CFC can be made if the phase information and the amplitude information are taken from two different electrodes, if neither electrode shows comparably strong CFC by itself. Such a finding will be in agreement with the dominant view that synchronization between brain oscillations reflects information integration (see for example Lakatos et al., 2005; Canolty and Knight, 2010; Sadeh et al., 2014).

Phase-amplitude cross frequency coupling is likely an important mechanism for synchronizing neural activity (3,11,20,28,29). Demonstrations of CFC have been described which correlate with cognitive function in several domains (2). The findings presented here do not suggest that CFC analysis is flawed, nor that current findings of CFC are necessarily misinterpreted. Rather, they suggest that there are multiple ways in which CFC can be generated, and call for careful consideration of parsimonious explanations when CFC is reported in a signal.

## Acknowledgments

The authors would like to thank Stephanie K. Ries from the Knight lab for permission to use ECoG data from her experiment. This study was funded by NINDS Grant R37 NS021135-27 to RTK, and the US-Israel Binational Research Foundation Grant 2013070 to RTK and LYD.

### Author Contributions

Conceptualization, EMG, BS, LYD and RTK; Methodology, EMG and BS; Software, EMG, BS and AW; Analysis, EMG and BS; Writing – Original Draft, EMG and BS; Writing – Review and Editing, RTK and LYD; Supervision, RTK and LYD; Funding Acquisition, RTK and LYD.

### Conflict of interest

None

